# Functional Ensemble Survival Tree: Dynamic Prediction of Alzheimer’s Disease Progression Accommodating Multiple Time-Varying Covariates

**DOI:** 10.1101/2020.02.17.952994

**Authors:** By Shu Jiang, Yijun Xie, Graham A. Colditz

## Abstract

With the exponential growth in data collection, multiple time-varying biomarkers are commonly encountered in clinical studies, along with rich set of baseline covariates. This paper is motivated by addressing a critical issue in the field of Alzheimer’s disease (AD) in which we aim to predict the time for AD conversion in people with mild cognitive impairment to inform prevention and early treatment decisions. Conventional joint models of biomarker trajectory with time-to-event data rely heavily on model assumptions and may not be applicable when the number of covariates is large. This thus motivated us to consider a functional ensemble survival tree framework to characterize the joint effects of both functional and baseline covariates in predicting disease progression. The proposed framework incorporates multivariate functional principal component analysis to characterize the changing patterns of multiple time-varying neurocognitive biomarker trajectories and then nest these features within an ensemble survival tree in predicting the progression of AD. We provide a fast implementation of the algorithm that accommodates personalized dynamic prediction that can be updated as new observations are gathered to reflect the patient’s latest prognosis. The algorithm is empirically shown to perform well in simulation studies and is illustrated through the analysis of data from the Alzheimer’s Disease Neuroimaging Initiative (ADNI). We provide implementation of our proposed method in `R` package `funest`.

## 1. Introduction

Alzheimer’s disease (AD) is one of the most prevalent disease worldwide which leads to memory loss and dementia [Mattson, 2004, LaFerla et al., 2007, Rabin et al., 2019]. Early detection is critical due to the lack of disease-modifying agents for patients diagnosed with AD. Mild cognitive impairment (MCI) is defined as the transition stage between the clinically normal and dementia state where it involves memory and language loss that is considered greater than expected age-related changes [Mattson, 2004]. As a result, MCI patients are typically enrolled as the target population for early prognosis and evaluation of therapies trials [Ewers et al., 2012]. There is considerable interest in identifying biomarkers or combination of covariates, so that the likelihood of predicting the neurodegenerative pathology due to Alzheimer’s disease for patients diagnosed with MCI can be greater. See Park et al. [2012], Ewers et al. [2012], Gomar et al. [2014] for example. Accurate and robust prediction of disease progression to AD is thus important and critical to move the field forward [Risacher et al., 2009].

Tremendous amounts of data are being collected in the hopes of finding significant factors that may be associated with AD progression. In the dataset that motivated this work, the Alzheimer’s Disease Neuroimaging Initiative (ADNI), the focus was on the collection of longitudinal assessments, magnetic resonance imaging and positron emission tomography imaging measures, as well as other biomarkers from blood and cerebrospinal fluid [Cuingnet et al., 2011]. Of those covariates collected in the cohort, many are time-varying. For example, the cognitive change in preclinical AD is a series of cognitive tests which are measured at each patient visits. The potential for discovery would be much greater in incorporating all available patient-specific covariates in predicting the progression of AD. However, the challenges may arise from i) high dimensionality of the baseline covariates; ii) presence of multivariate time-varying biomarkers; iii) non-linear and complex relationship between the covariates and the time-to-event outcome. A natural question then is how to best utilize this information to improve prediction performance to inform prevention and early treatment decisions.

Existing methods in the literature, such as the joint model, hinges on the pre-specified model assumptions for both the time-varying biomarker and the survival outcome [Rizopoulos, 2012]. However the nature of time-varying biomarkers may vary under different clinical settings, making it difficult to identify a suitable model. For illustration purposes, we present in Figure 1, the raw longitudinal trajectories for two of the longitudinal cognitive measures for 50 randomly selected MCI patients in ADNI. We can see that both the Mini Mental State Examination (MMSE, left) and Functional Activities Questionnaire (FAQ, right) trajectories have changing patterns over time and are highly variable within and between patients. In addressing this concern, nonparametric methods such as splines or kernel smoothing, have been adopted in the literature for prediction using the denoised smoothed values of the biomarker trajectories [Wu and Chiang, 2000, Welsh et al., 2002]. More recently, functional approaches such as functional principal component analysis (FPCA), has become a popular alternative for modeling time-varying predictors due to its ability to use extracted features (changing patterns) in addition to the denoised smoothed values which will likely improve prediction [Ramsay and Silverman, 2004, Wang et al., 2016]. Examples of functional data analysis applied to time-to-event data include Yan et al. [2017, 2018], Kong et al. [2018]. However all of the aforementioned methods focuses on the dynamic prediction of time-to-event outcome with a single time-varying biomarker. As a result, Li and Luo [2019] recently proposed the use of multiple longitudinal biomarkers in predicting the disease progression. However, their method contingent on the proportional hazards model which may not be realistic and viable especially when the number of covariates is large.

**Fig 1.**
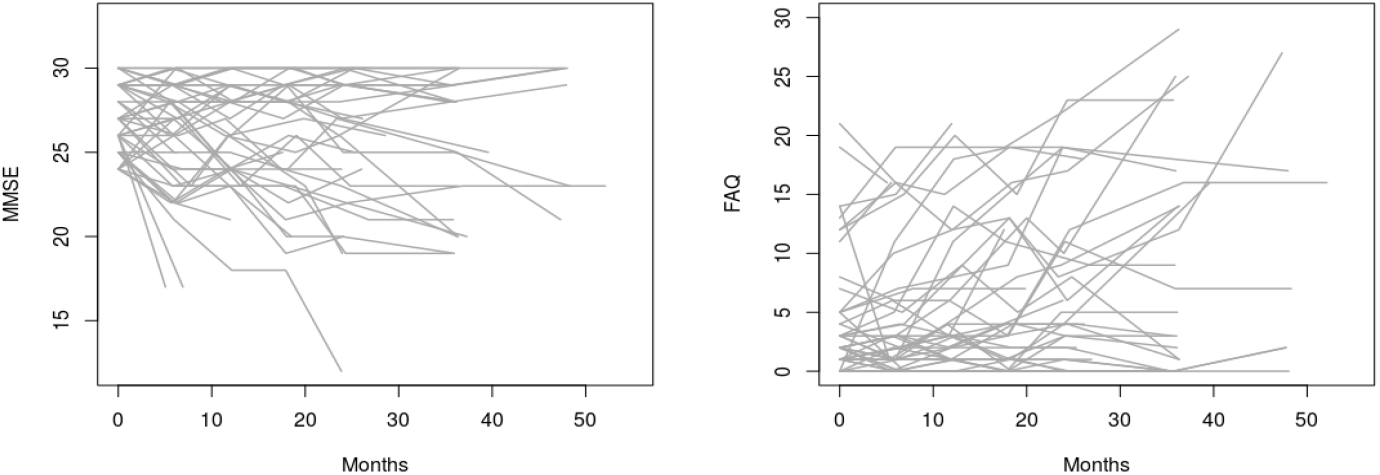
Longitudinal trajectories of Mini Mental State Examination (MMSE, left) and Functional Activities Questionnaire (FAQ, right) of 50 randomly selected MCI patients in ADNI.

In this article, we propose a unified strategy for dynamic prediction that does not depend on the model specification and can handle high dimensional baseline and multivariate time-varying covariates in the presence of right censoring. The proposed approach is entirely data-driven and can be stated in terms of three main steps. (1) First, extract features from the multivariate time-varying covariates such that the changing patterns can be summarized by a set of functional basis functions and the associated individualized functional scores. (2) Then, construct candidate estimators based on the extracted features and observed data. (3) Last, apply cross-validation to select the optimal estimator among all candidates in step 2. Specifically, we adopt tree-based methods in this paper where the possible candidate estimators in step 2 are generated by repeated binary recursive partitions [Ishwaran et al., 2008, 2011]. Tree-based methods facilitate a comprehensive modeling scheme and are appealing for their ability to handle data with high-dimensional covariates, facilitate complex and nonlinear relationship between predictors and outcomes and relax the proportional hazard assumption [Taylor, 2011, Jiang, 2019]. Given the tree-based estimators in step 2, the optimal estimator in step 3 can be selected via cross-validation by tuning the number of functional basis functions from step 1 and tree-based parameters which we discuss in detail in Section 2. The proposed method will reinforce model robustness and prediction accuracy and serve as a valuable tool for researchers in conducting future research.

The remainder of this paper is organized as follows. In Section 2, we define notation and describe the model setup. In particular, we give detailed discussion on the multivariate principal component analysis (MFPCA) for feature extraction from multiple time-varying covariates and the construction of the functional ensemble survival tree for conducting individualized dynamic prediction. We investigate the finite sample performance in intensive simulation studies in Section 3 and provide publicly available code in `R` package `funest`. An application involving ADNI is given in Section 4, and concluding remarks and topics for future research are given in Section 5.

## 2. Notation and Method

### 2.1. Functional Ensemble Survival Tree

Random survival forest (RSF) is an ensemble tree method that has been widely adopted for the analysis of right-censored survival data. The goal of constructing the RSF is to train a model that learns from the available functional and baseline covariates in the cohort, such that the model can be used to make risk predictions for new patients conditioning on partially observed data. The focus of this subsection is on model construction and we elaborate on individualized dynamic prediction in Section 2.2.

Typical RSF can not take longitudinal covariates directly as inputs. To extend the survival tree on the basis of longitudinal covariates, we first characterize the changing patterns of the time-varying biomarkers via MFPCA. We start by setting up the functional framework for single time-varying biomarkers and then expand to the multivariate setting. We let 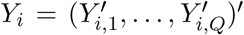 be the observed time-varying biomarkers for individual *i*, *i* = 1, …, *n*. The *q*th time-varying biomarker is denoted by 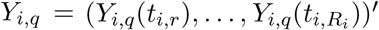 where *R*_*i*_ reflects random and irregular individual-specific visits, *q* = 1, …, *Q*. We assume that the *q*th observed trajectory, ∀*q* ∈ {1, …, *Q*}, is recorded with error,

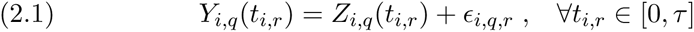

where *Z*_*i,q*_(*t*_*i,r*_) denotes the denoised mean value of *Y*_*i,q*_(*t*_*i,r*_) for *t*_*i,r*_ ∈ [0, *τ*] and *τ* denotes the maximum follow up time in the cohort. The error term is assumed to have *E*(*ϵ*_*i,q,r*_) = 0 and 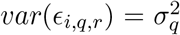 where *t*_*i,r*_, *Z*_*i,q*_ and *ϵ*_*i,q,r*_ are assumed to be mutually independent [Yao et al., 2005].

Under the functional framework, we assume that *Z*_*i,q*_ = {*Z*_*i,q*_(*t*), ∀*t* ∈ [0, *τ*]} are realizations of a stochastic process *Z*_*q*_(*t*) in a square integrable functional space with domain *τ*. The stochastic process is assumed to have mean function *E*[*Z*_*q*_(*t*)] = *µ*_*q*_(*t*) and covariance operator *C*_*q*_(*t, s*) = *Cov*(*Z*_*q*_(*t*), *Z*_*q*_(*s*)) for ∀*t*, *s* ∈ [0, *τ*]. Then by Mercer’s theorem [Mercer, 1909],

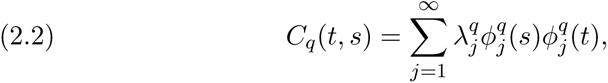

where 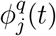 is the *j*th orthonormal eigenfunction and 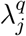 is the corresponding eigenvalue where 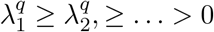, *j* = 1, …, *∞*. This decomposition thus allows us to characterize each functional observation *Z*_*iq*_(*t*) as

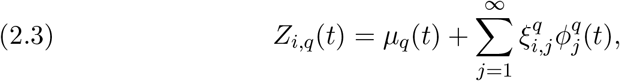

where 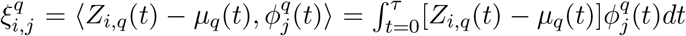, is the *j*th functional principal component (FPC) score for individual *i*. According to the Karhunen–Loève theorem [Ramsay and Silverman, 2004], each curve *Z*_*i,q*_(*t*), ∀*t* ∈ [0, *τ*], can then be characterized by the infinite sequence of FPC scores 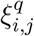, *j* = 1, …, *∞*. In practice, an approximation of (2.3) is usually carried out by truncating the infinite summation to the first *M*_*q*_ terms where *M*_*q*_ could be determined by, for example, Akaike information criterion (AIC) or the total variance explained (TVE) [Wang et al., 2016]. For estimation, given the observed data, we adopt the Principal Analysis by Conditional Estimation (PACE) algorithm for its well-known property of accommodating sparse longitudinal observations as is the case in our motivating study [Yao et al., 2005]. Specifically, we use PACE algorithm to facilitate the estimation of the discretized mean function 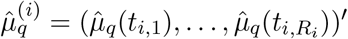, the *R*_*i*_ × *R*_*i*_ empirical covariance matrix 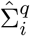 and the corresponding eigenvectors 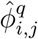 and eigenvalues 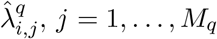. Then the univariate FPC scores for the *q*th biomarker trajectory for *i*th individual can be estimated as

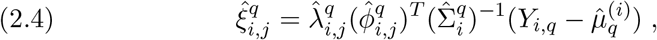

*j* = 1, …, *M*_*q*_, for *q* = 1 …, *Q*.

Next we combine the *Q* univariate time-varying biomarkers via MFPCA following Happ and Greven [2018]. We let 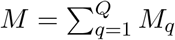 and 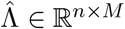 be an *n* × *M* matrix for which the *i*th row is 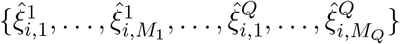. In the multivariate setting we aim to perform a matrix eigenanalysis such that we can estimate the corresponding eigenvectors 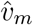, from the empirical block matrix 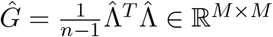, *m* = 1, …, *M*. Note that MFPCA indirectly accommodates the potential correlations among multiple trajectories via correlation among the FPC scores by pooling all estimated eigenvectors from the univariate biomarkers in the block matrix 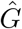. The eigenvectors 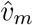 thus contain the information of correlations across different time-varying biomarkers. As a result, the multivariate eigenfunctions are estimated as

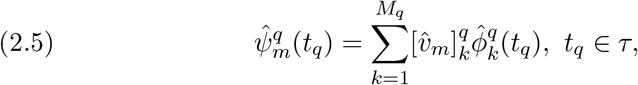

where 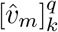 denotes the *k*th entry in the *q*th block of 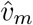, *q* = 1, …, *Q*, *m* = 1, …, *M*. The corresponding individual-specific MFPC scores can thus be estimated as

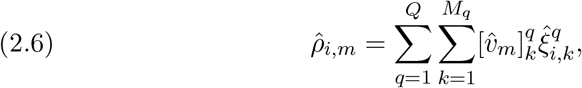

*m* = 1, …, *M*. Similar to the univariate setting, the optimal number of MFPCs, {*D*: *D* ≤ *M*}, can be chosen based on, for example, TVE or AIC.

The RSF can be easily constructed once the MFPCA scores have been estimated as in (2.6). Within the forest, every tree in the forest is grown from a single node to a tree with multiple terminal nodes. Specifically, each decision tree is grown by partitioning individuals at each node into two groups, where the split is chosen under a user-specified splitting rule. Node splitting rules often are determined with the goal to either maximize within-node homogeneity or between-node heterogeneity. The standard split criterion for survival trees is the log-rank statistic to maximize the survival differences at each node which has been widely used and implemented [Ishwaran et al., 2011]. Other splitting criterion such as the maximally selected rank statistic has been recently developed for its well-known unbiased split variable selection property [Wright and Ziegler, 2015]. In each terminal node of a tree, the survival function is estimated using the Kaplan–Meier estimator, utilizing only the observations from the same terminal node. Note that several parameters needs to be tuned via cross-validation. In particular, the prediction error needs to be assessed with, for example, various number of trees, number of covariates to split on and the minimal terminal node size. In addition, we may also tune the number for MFPCs that are nested within the RSF for a better prediction performance. See Section 3 for more details.

### 2.2. Individualized Dynamic Prediction

We let *n* denote the number of individuals in the training cohort and *n* + 1 be the new individual who is event-free and has observation up to some time *t*^*^, *t*^*^ < *τ*. For each single tree *b*, *b* = 1, …, *B*, prediction of the survival probability at *t*^*^ + *Δt* < *τ*, is made by dropping the new individual *n* + 1’s observations down the tree as

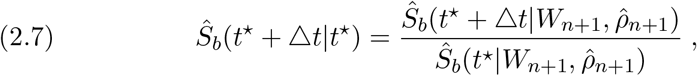

where *W*_*n*+1_ is the baseline covariates for individual *n*+1 of dimension *P* × 1. The MFPC scores 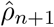 can be obtained by first estimating the univariate FPC scores from (2.4),

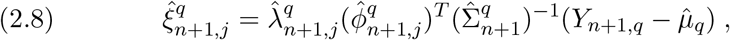

*j* = 1, …, *M*_*q*_, *q* = 1, …, *Q*. We then pass these FPC scores to (2.6) to obtain the MFPCA scores 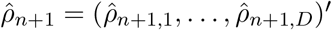. The final prediction from the forest is estimated by averaging over *B* trees in as

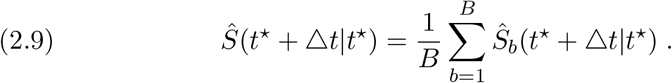

## 3. Simulation Study

We conduct intensive simulation studies to investigate the finite sample performance of our proposed method in this section. We aim to mimic the motivating application and simulate *n* = 400 individuals in each dataset with *nsim* = 500. The individual-specific visit times {*t*_*i,r*_, *r* = 1, 2, …, 7} are generated from the Gaussian distribution centered at 0, 3, 6, 9, 12, 15, and 18 with standard deviation of 0.1 except the initial baseline visit which is fixed at 0.

We assume that the time-varying biomarkers are recorded with error, *Y*_*i,q*_(*t*_*i,r*_) = *Z*_*i,q*_(*t*_*i,r*_)+*ϵ*_*i,r,q*_, where *ϵ*_*i,r,q*_ ∼ *N* (0, 1) and *q* = 1, 2, 3. We consider both the linear and non-linear longitudinal trajectories in a similar fashion as Li and Luo [2019]. Specifically in the linear setting, we simulate

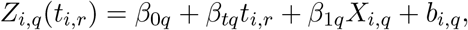

where [*β*_01_, *β*_02_, *β*_03_] = [1.5, 2, 0.5], [*β*_*t*1_, *β*_*t*2_, *β*_*t*3_] = [1.5, −1, 0.6], and [*β*_11_, *β*_12_, *β*_13_] = [2, −1, 1]. We simulate *X*_*i,q*_ ∼ *N* (3, 1) for *q* = 1, 2, 3 and the individual-specific random effects [*b*_*i*,1_, *b*_*i*,2_, *b*_*i*,3_] ∼ *MV N* (**0**, Σ) with

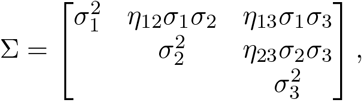

where 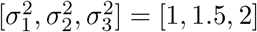, and [*η*_12_, *η*_13_, *η*_23_] = [−0.2, 0.1, −0.3].

The nonlinear trajectories for each individual *i* is assumed to follow a piecewise model

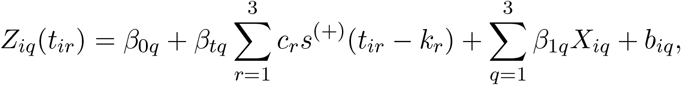

where [*c*_1_, *c*_2_, *c*_3_] = [1.2, 0.7, 0.5], [*k*_1_, *k*_2_, *k*_3_] = [0, 6, 13], and

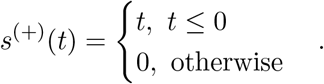

We assume a porportional hazards model in this simulation where the conditional hazard function follows

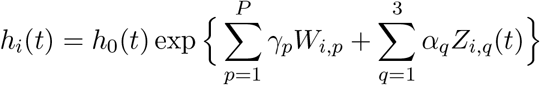

where *α*_*q*_ is set to be (0.1, −0.1, 0.2) for *q* = 1, 2, 3 respectively. We consider four different scenarios for the set of fixed covariates *W*_*i*_ = (*W*_*i*,1_, …, *W*_*i,P*_)^*t*^. In the first two scenarios we set *ρ* = 0.2 and 0.5 for *P* = 20, 100 respectively to represent strong autoregressive dependence where *W*_*i*_ ~ MVN(0, Σ^(*W*)^) with the (*k, l*)th component of Σ^(*W*)^ define as

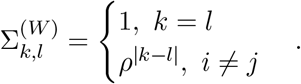

In the last two scenarios, we consider binary covariates with *P* (*W*_*i,p*_ = 1) = 0.5 similarly under *ρ* = 0.2 and 0.5 for *P* = 20, 100. We set the associated coefficients *γ*_*p*_ = (−2.5, −0.5, −0.15, −0.15, −0.1) for *p* = 1, … 5 so that high values of *γ*_1_, … *γ*_5_ are associated with shorter times to the event, and *γ*_*p*_ = 0 for *p* = 6, …, *P*. The elements of *W*_*i*_ with non-zero coefficients were chosen to give both weak and strong dependence within the set of important covariates accompanied with set of noise variables. With the above setups, we are then ready to simulate 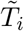 and *C*_*i*_. As demonstrated in Austin [2012] the survival time *T*_*i*_ can be generated from the inverse of the cumulative hazard function 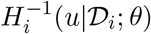 where *H*(*t*) is the cumulative hazard function and *u* ∼ unif(0, 1). We have simulated under the independent censoring scheme, where the censoring time is set to follow a uniform distribution *unif* (0, *C*_*m*_) where *C*_*m*_ is set such that the % of being censored by the end of the study is 30%.

In the simulations that we conducted, 300 individuals were randomly chosen from each simulated dataset to train the model and the 100 used to evaluate the prediction performance. To avoid overfitting, we employed a 5-fold inner and outer cross-validation. Specifically for the inner cross-validation, an optimal ensemble survival tree model was built and selected based on the best prediction performance by tuning the parameters in each fold. For each fold in the outer cross-validation, the prediction accuray measure is recorded dynamically for each time window (*t*^*^, *t*^*^ + *Δt*] conditional on data observed up to *t*^*^,*t*^*^ = 6, 9, forecasting *t*^*^ + *Δt* for *Δt* = 3, 6.

Table 1 illustrates simulation results from the nonlinear setting. Additional simulation results under the linear setting are provided in Table 2 within the Supplemental Material for interested readers. As shown in Table 1, the AUC [Li et al., 2015] outputted from the proposed method are in good agreement with the true AUC confirming a satisfactory model discrimination. Additionally, we see that the Brier scores [Schoop et al., 2008] are very close to zero which confirms good model calibration as well. From our results, we can see that when the signal-to-noise ratio (S:N) decreases from 5: 15 to 5: 95, the proposed model retains robust performance in both the AUC and Brier score.

**Table 1.**
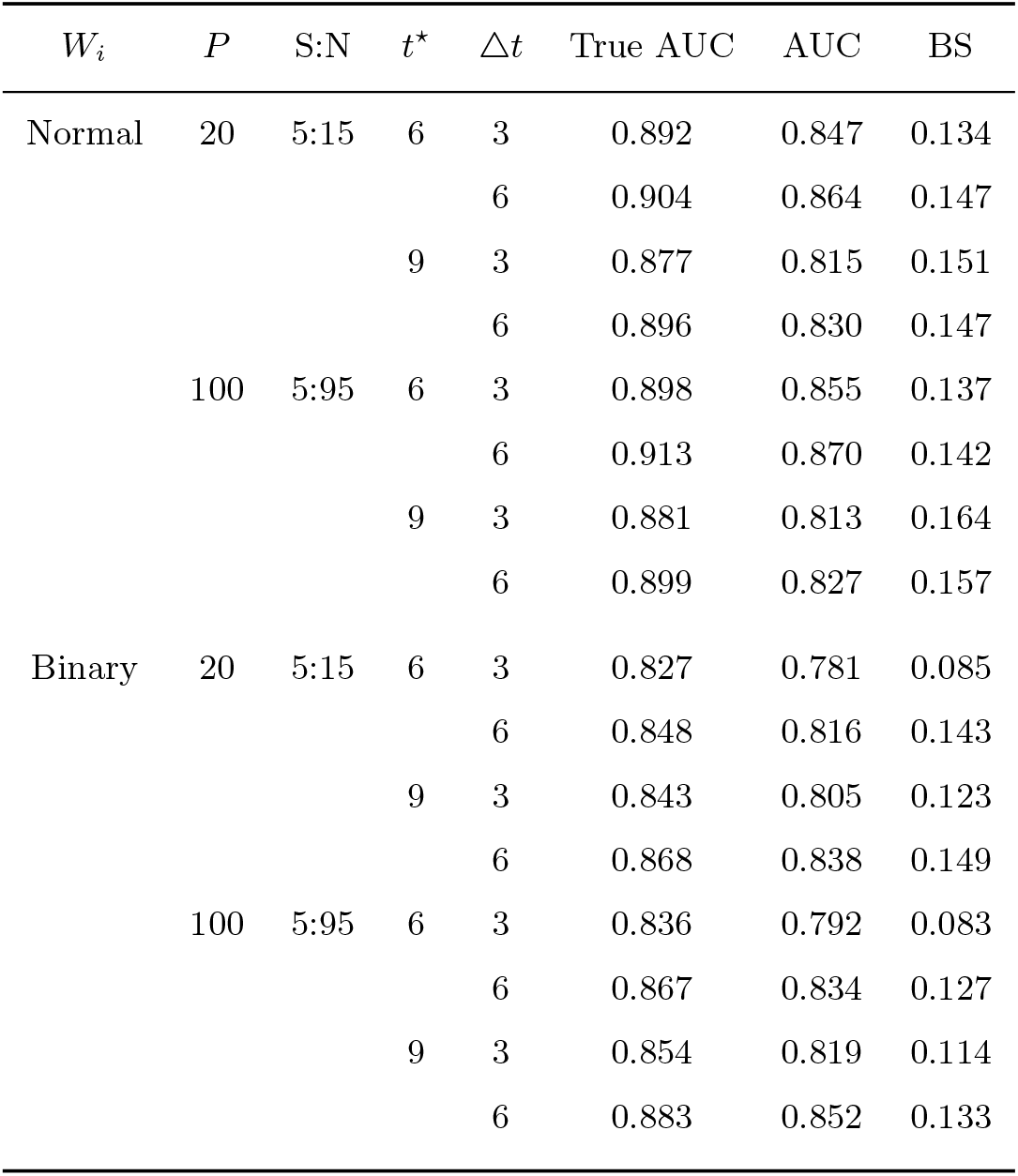
Estimated AUC(t^*^, t^*^ + Δt) and Brier score(t^*^, t^*^ + Δt) under the nonlinear setting via functional ensemble survival tree; n = 400, nsim = 500, S:N = signal-to-noise ratio.

**Table 2.**
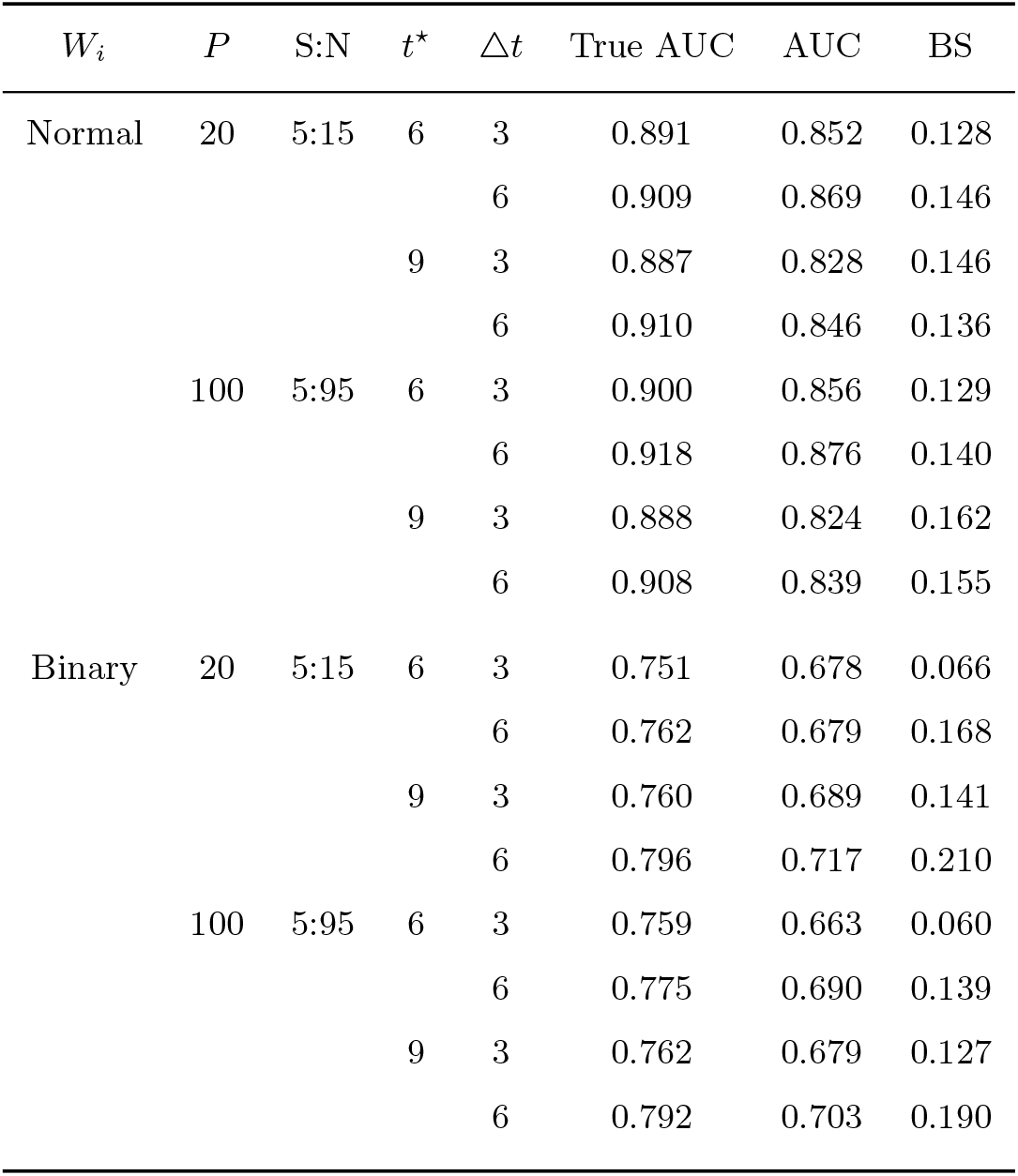
Estimated AUC(t^*^, t^*^ + Δt) and Brier score (t^*^, t^*^ + Δt) under the linear setting via functional ensemble survival tree; n = 400, nsim = 500, S:N = signal-to-noise ratio.

The proposed method has been implemented in our `funest` package which utilizes the well developed `ranger` that wraps the implementation of random forest in C++. The computational speed of our package is outstanding, as it takes less than 30 seconds for growing and making dynamic predictions on the functional ensemble survival forest with *∼*2500 trees on a desktop with i7-7700 CPU. In addition, the package naturally takes advantage of the multi-core processor when running in a larger scale computational environment which warrants a even more promising computational speed.

## 4. Alzheimer’s Disease Neuroimaging Initiative

The data used in this section were obtained from the Alzheimers Disease Neuroimaging Initiative (ADNI) database^1^. We treat the conversion from MCI to AD as the time-to-event outcome and focus on 317 patients who have been diagnosed with MCI in ADNI-1. Out of those who were diagnosed with MCI, 141 of them progressed to AD before the end of the study. Patients were assessed at baseline, 6, 12, 18, 24, and 36 months in ADNI-1 with additional annual follow-ups in ADNI-2 resulting in an average follow-up period of 33.4 (sd = 14.1) months. The corresponding average number of visits recorded was of 6.3 (sd = 2.3). Table 3 in the Supplemental Material shows the list of variables that we consider in this section. We have focused on five time-varying neurocognitive markers as well as other baseline covariates that have been well studied in the Alzheimer’s literature [Mattson, 2004, LaFerla et al., 2007, Gomar et al., 2014].

**Table 3.** Covariates used in ADNI dataset; ^†^ represents a time-varying covariate.

| Variable | Description |
| --- | --- |
| ADAS-Cog 13<sup>†</sup> | Alzheimer Disease Assessment Scale-Cognitive 13 items |
| RAVLT.immediate<sup>†</sup> | Rey Auditory Verbal Learning Tests immediate score |
| RAVLT.learning<sup>†</sup> | Rey Auditory Verbal Learning Tests learning score |
| FAQ<sup>†</sup> | Functional Assessment Questionnaire |
| MMSE<sup>†</sup> | Mini Mental State Examination |
| mPACCdigit.bl | baseline preclinical Alzheimer's cognitive composite score |
| MidTemp.bl | baseline Middle temporal gyrus volume |
| Hippocampus.bl | baseline Hippocampus gyrus volume |
| Fusiform.bl | baseline Fusiform gyrus volume |
| Entorhinal.bl | baseline Entorhinal volume |
| Ventricles.bl | baseline Ventricles volume |
| ADAS-Cog13.bl | baseline Alzheimer Disease Assessment Scale-Cognitive 13 items |
| RAVLT.immediate.bl | baseline Rey Auditory Verbal Learning Tests immediate score |
| RAVLT.learning.bl | baseline Rey Auditory Verbal Learning Tests learning score |
| FAQ.bl | baseline Functional Assessment Questionnaire |
| MMSE.bl | baseline Mini Mental State Examination |
| APOE4 | apolipoprotein E gene indicator |
| AGE | age at recruitment |
| PTGENDER | gender |
| PTEDUCAT | years of education |

Figure 2 illustrates the dynamic prediction performances under different models. In particular, we considered model 1 which is the full model that includes all available covariates and model 2 which includes only baseline covariates (including baseline measures for the time-varying markers). To avoid overfitting, we employed a 5-fold inner and outer cross-validation in this analysis similar to our simulation study. An optimal ensemble survival tree model was built and selected based on the best prediction performance by tuning the parameters in each fold in the inner cross-validation. For each fold in the outer cross-validation, the prediction accuracy measure is recorded dynamically for each time window (*t*^*^, *t*^*^ + Δ*t*] conditional on data observed up to *t*^*^,*t*^*^ = 6, 12, 18, 24 (month), forecasting *t*^*^ + Δ*t* for Δ*t* = 6 month. It is apparent that model 1 achieves a better AUC(*t*^*^, *t*^*^ + Δ*t*) and BS(*t*^*^, *t*^*^ + Δ*t*) dynamically over all time points. This suggests that the inclusion of the changing pattern of time-varying covariates indeed facilitates a better model discrimination and calibration.

**Fig 2.**
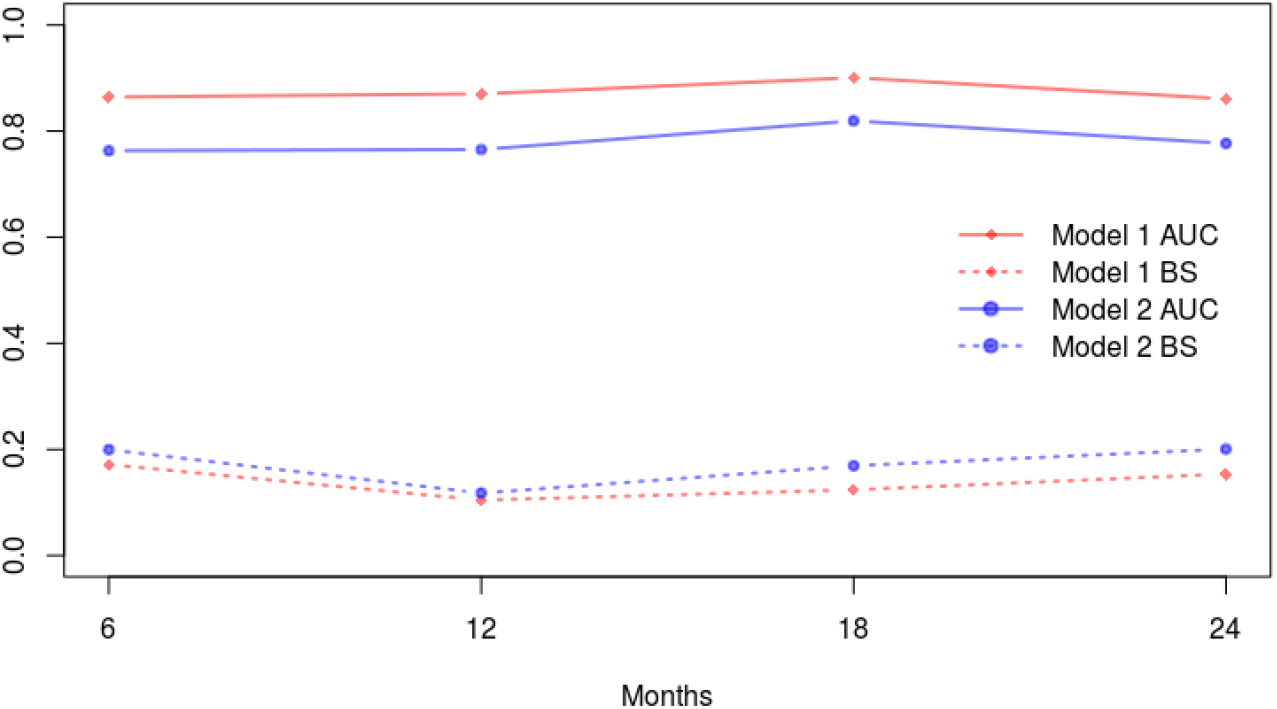
Comparison of dynamic prediction performances of the full model (model 1) and baseline model (model 2) under a 5-fold cross-validation. AUC(t^*^, t^*^ +6) and BS (t^*^, t^*^ +6) conditions on data observed prior to t^*^ = 6, 12, 18, 24 (month) in forecasting t^*^ + 6 under a sliding window framework.

We further illustrate the variable importance ranking via the variable permutation importance measure as shown in Figure 3. A variable is identified as important if it exerts a positive effect on the prediction performance. A greater value of permutation measure on a variable implies that the variable is more important for the overall predictive accuracy; see Nembrini et al. [2018] for more details. As a result, we see from Figure 3 that the first principal component (PC1) stands out with significantly large permutation importance measure in relation to other variables. This finding suggests that the contribution of the changing patterns of time-varying covariates in predicting the progression of AD for those who are diagnosed as MCI is much greater relative to the fixed baseline covariates. Such finding is indeed in agreement with what we observe in Figure 2.

**Fig 3.**
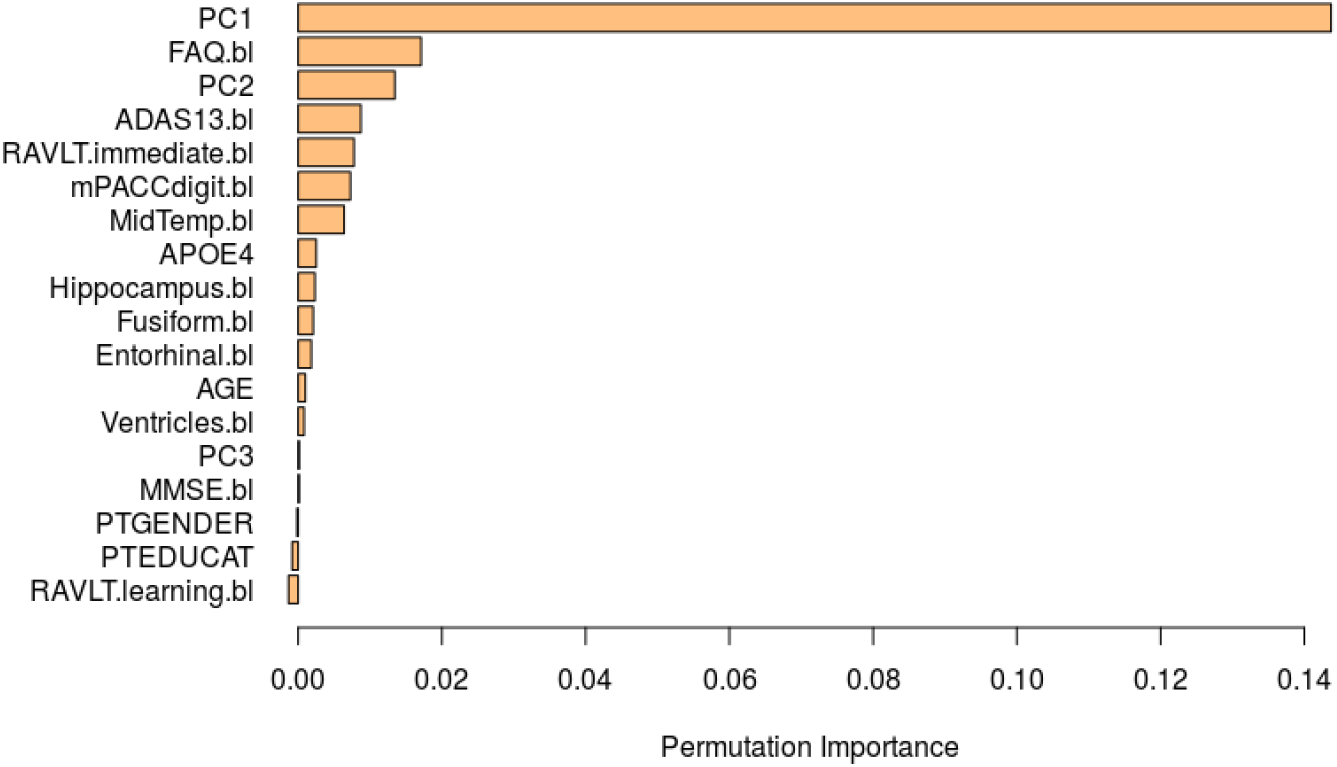
Varbiable permutation importance barplot.

Lastly, we demonstrate individualized dynamic prediction where we randomly set aside a single patient from the cohort. The model was trained on the remainder of the cohort leaving this single patient out, such that we are able to visualize the predicted future biomarker trajectory and risk conditional on partial profile. Figure 4 shows two of the five neurocognitive markers (i.e., ADAS-COG13 and FAQ) that we use in the model for ease of visualization. The dashed line in Figure 4 represents the last time the biomarker has been recorded for the patient. From the first column of Figure 4, we can see that the predicted trajectories of the ADAS-COG13 and FAQ are in great harmony with the true values. Correspondingly, the predicted AD-free probability for the patient is shown as a function of time in the second column where splines have been adopted to provide a smooth curve. The predicted low risk should be consistent with what one would expect from the stability of neurocognitive marker measurements. A full set of neurocognitive markers and their associated predictions are provided in Figure 5 in Supplemental Material for interested readers.

**Fig 4.**
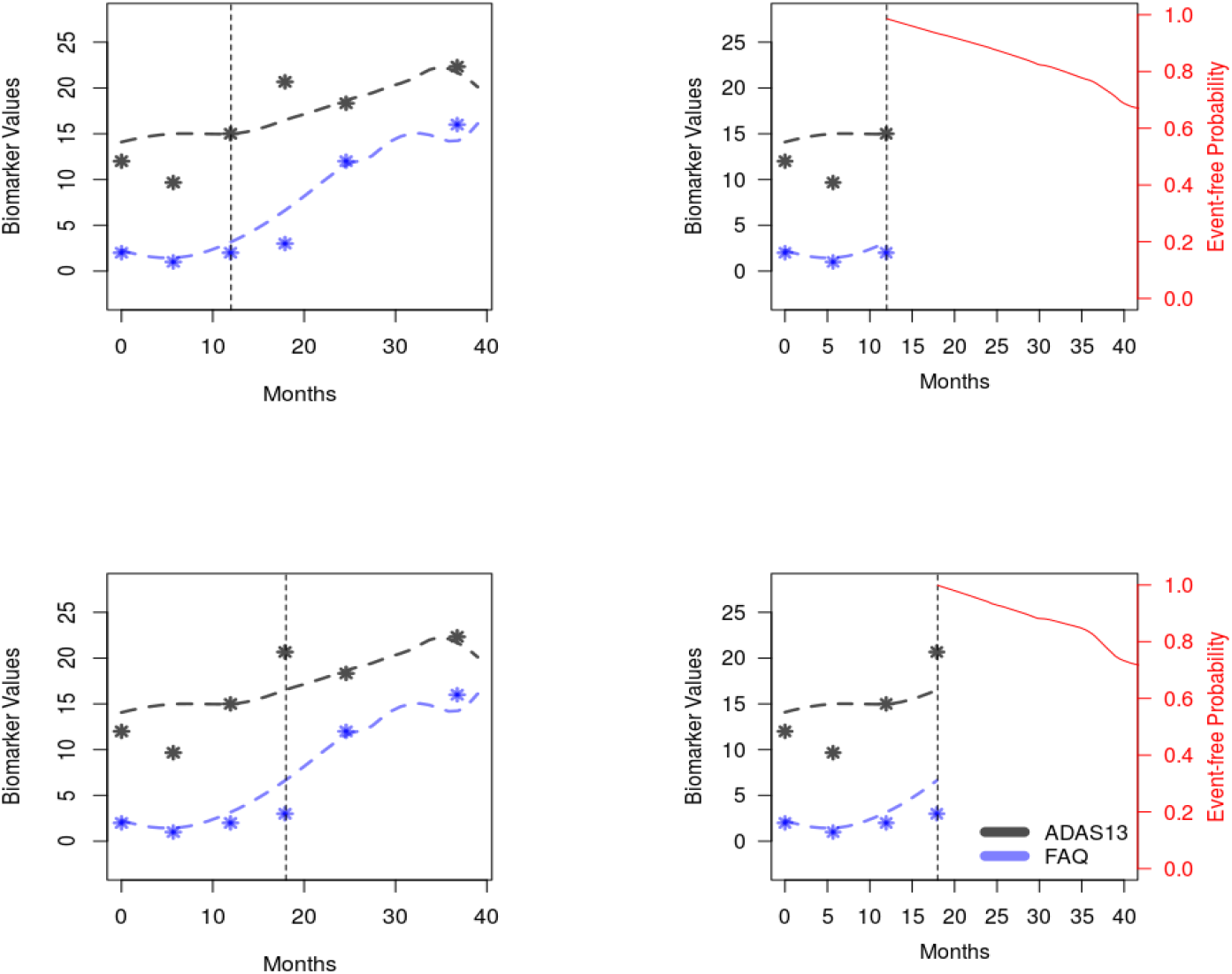
Predicted trajectories of ADAS-COG13 and FAQ in the first column and predicted AD-free probability in the second column conditional on partially observed marker values prior to the dashed line; dashed line represents the last time the biomarker has been recorded for the patient.

**Fig 5.**
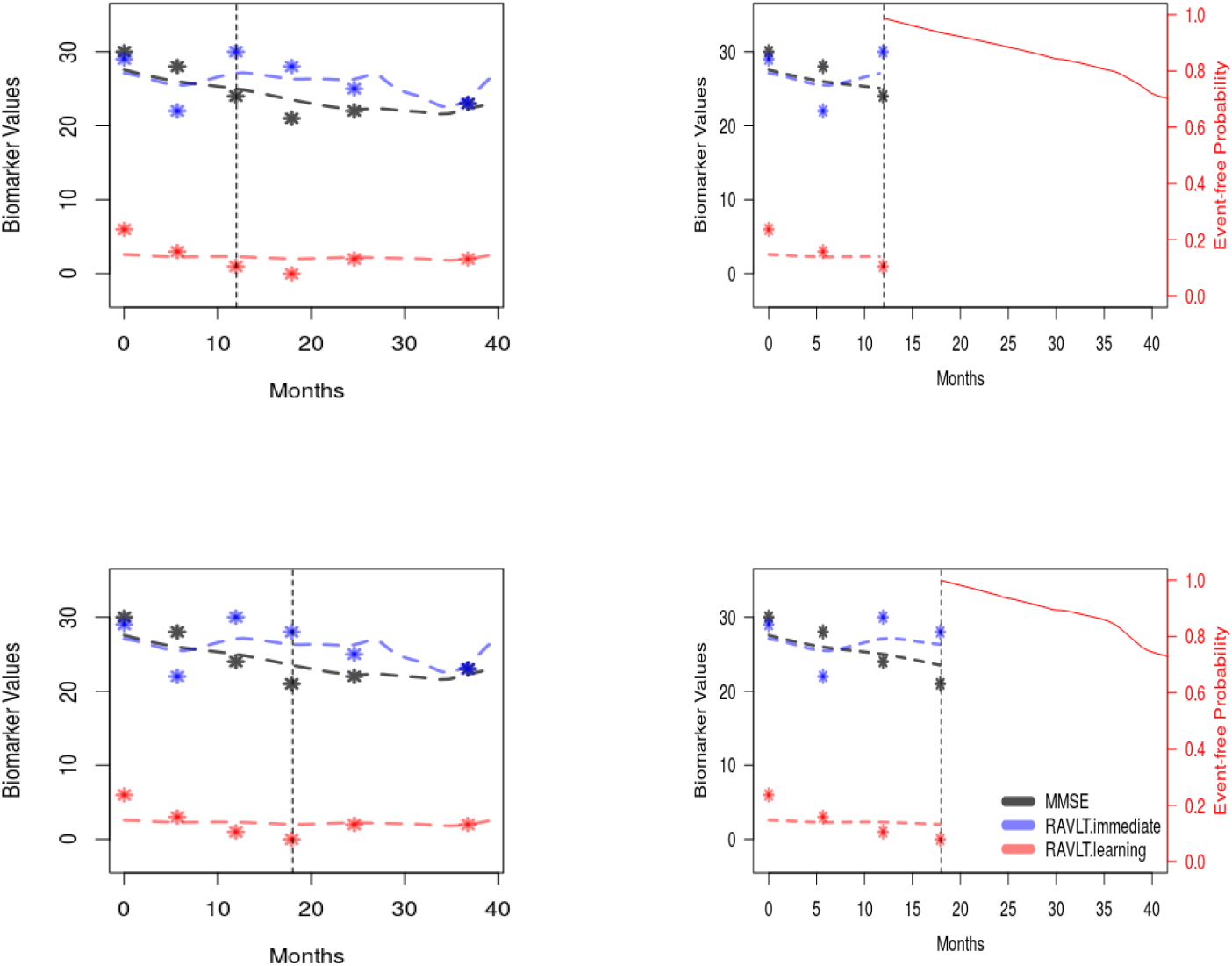
Predicted trajectories of MMSE, RAVLT.immediate and RAVLT.learning in the first column and predicted AD-free probability in the second column conditional on partially observed marker values prior to the dashed line; dashed line represents the last time the biomarker has been recorded for the patient.

## 5. Discussion

We have formulated the functional ensemble survival tree framework that facilitates individualized dynamic prediction for disease progression accommodating multiple time-varying biomarkers. The proposed framework is fully data-driven and therefore removes the burden of the need to impose model assumptions on both the time-varying trajectories and the survival distribution. Specifically, we adopt MFPCA to characterize the changing pattern of the multivariate time-varying biomarkers which effectively captures the correlation among them. We also nest these extracted features into the ensemble survival tree which accommodates dynamic prediction under high dimensionality of baseline covariates and complex associations between the covariates and the time-to-event outcome. We investigate the empirical performance of the proposed algorithm and show that the model is robust and has a good discrimination and calibration via both the AUC and Brier score. We describe how to conduct individualized dynamic prediction and illustrate the proposed framework in the ADNI dataset. Furthermore, we make the proposed algorithm publicly available in `R` package `funest`. This could help physicians to predict future course of the biomarker trajectories as well as the associated risk of AD which in turn, could facilitate identifying high risk individuals for prevention trials and treatment interventions.

A limitation in all tree-based methods is the lack of interpretability. However, in analyzing the ADNI dataset, we provided the variable permutation importance measure which identifies variable that are important contributors for the overall predictive accuracy. Our findings from ADNI suggest that time-varying trajectories play a major role in predicting AD progression for those that are diagnosed with MCI. We did not use the genetic marker data in our analysis as ADNI is an ongoing project that currently only has a sparse number of individuals who have their genetic profiles available. However, the model set up and the software distributed in this article warrants further research incorporating a richer set of covariates.

## Acknowledgements

This work is supported in part by funding from the Foundation for Barnes Jewish Hospital and P30 CA091842. Data used in preparation of this article were obtained from the Alzheimers Disease Neuroimaging Initiative (ADNI) database (http://adni.loni.usc.edu). As such, the investigators within the ADNI contributed to the design and implementation of ADNI and/or provided data but did not participate in analysis or writing of this report. A complete listing of ADNI investigators can be found at: http://adni.loni.usc.edu/wpcontent/uploads/howtoapply/ADNIAcknowledgementList.pdf.

## SUPPLEMENTARY MATERIAL

### Additional Simulation Results

The prediction performance measures under the linear setting in Section 3 are displayed in Table 2.

### Additional Real Data Results

Table 3 gives a full list of the covariates that we have used in Section 4 for the ADNI dataset. Additional individualized dynamic prediction plots with all neurocognitive markers are displayed in Figure 5.

().

## Footnotes

1 http://adni.loni.usc.edu/

